# Incorporating evaporative water loss into bioenergetic models of hibernation to test for relative influence of host and pathogen traits on white-nose syndrome

**DOI:** 10.1101/750257

**Authors:** Catherine G. Haase, Nathan W. Fuller, C. Reed Hranac, David T. S. Hayman, Liam P. McGuire, Kaleigh J. O. Norquay, Kirk A. Silas, Craig K. R. Willis, Raina K. Plowright, Sarah H. Olson

## Abstract

Hibernation consists of extended durations of torpor interrupted by periodic arousals. The ‘dehydration hypothesis’ proposes that hibernating mammals arouse to replenish water lost through evaporation during torpor. Arousals are energetically expensive, and increased arousal frequency can alter survival throughout hibernation. Yet we lack a means to assess the effect of evaporative water loss (EWL), determined by animal physiology and hibernation microclimate, on torpor bout duration and subsequent survival. White-nose syndrome (WNS), a devastating disease impacting hibernating bats, causes increased frequency of arousals during hibernation and EWL has been hypothesized to contribute to this increased arousal frequency. WNS is caused by a fungus, which grows well in humid hibernaculum environments and damages wing tissue important for water conservation. Here, we integrated the effect of EWL on torpor expression in a hibernation energetics model, including the effects of fungal infection, to determine the link between EWL and survival. We collected field data for *Myotis lucifugus*, a species that experiences high mortality from WNS, to gather parameters for the model. In saturating conditions we predicted healthy bats experience minimal mortality. Infected bats, however, suffer high fungal growth in highly saturated environments, leading to exhaustion of fat stores before spring. Our results suggest that host adaptation to humid environments leads to increased arousal frequency from infection, which drives mortality across hibernaculum conditions. Our modified hibernation model provides a tool to assess the interplay between host physiology, hibernaculum microclimate, and diseases such as WNS on winter survival.

---

In periods of food scarcity, hibernators conserve energy by entering torpor, during which body temperature (T_b_) is maintained near hibernaculum temperature and metabolic rate is lowered to reduce energy demands [1]. There are several hypotheses proposed to explain periodic arousals, but two of the most prominent are linked to water balance: 1) the dehydration hypothesis [2,3]; and 2) the need to excrete metabolic byproducts [4]. The dehydration hypothesis suggests that hibernators arouse periodically after a threshold of total body water is reached [5,6]. The metabolic byproducts hypothesis suggests that bats accumulate byproducts from biochemical reactions during torpor, and these byproducts need to be excreted as waste as they can be damaging to cellular function [4]. Both of these hypotheses are affected by microclimate, which is supported by empirical evidence of the relationship between hibernaculum temperature and relative humidity and torpor bout duration [3,5,7,8]. Hibernators do not normally defecate or urinate during torpor [9], thus water lost during inactive periods of hibernation is assumed to be from evaporation. Evaporative water loss (EWL) is comprised of respiratory and cutaneous water loss [10,11] and is driven by the difference in water vapor pressure between the surface of an animal and the surrounding air, which, in turn, is determined by the saturation of the air given air temperature [12]. The dehydration hypothesis is supported by correlations between torpor bout duration/arousal frequency and hibernaculum temperature and relative humidity in both free living and laboratory conditions [2,5,13].

Though arousals only make up a small portion of hibernation time, these periods account for the majority of the winter energy budget [1,14]. Therefore, the influence of microclimate on arousal frequency is critical for over-winter fat loss. However, the influence of microclimate has recently become an important question in the context of white-nose syndrome (WNS). WNS is a rapidly spreading infectious disease that has led to high mortality rates in hibernating bats across eastern and central North America. It has been proposed that increased EWL from infection could be a trigger of increased arousals associated with WNS [13,15].

The causal agent of WNS is a psychrophilic fungus, *Pseudogymnoascus destructans*, which erodes wing tissue [16,17]. Mechanistic models and empirical evidence connecting *P. destructans* infection to altered torpor-arousal cycles suggest that ulceration of the highly vascularized wing tissues causes increased fluid and water loss [13,18–21]. The growth rate of *P. destructans* is linked to both ambient temperature [22] and humidity [23], with higher fungal growth rates in environmental conditions frequently found in bat hibernacula.

Although studies have linked arousal frequency with survival, and hibernaculum microclimate and EWL with arousal frequency, none have explicitly considered the effect of EWL on survival to our knowledge. Hibernation energetic models are commonly used to understand energy consumption over winter but have yet to account for water balance and its effect on arousal behavior. With the current threat of WNS, a disease that potentially directly impacts water balance, it is important to understand the implications surrounding the association of EWL, energy consumption, and survival. We therefore developed a hibernation energetics model that incorporates water balance to assess the effects of the dehydration hypothesis on survival of *Myotis lucifugus*, a wide-ranging species that is heavily impacted by WNS. Using WNS as a study system, we tested the hypothesis that increased EWL from fungal infection results in greater energy consumption due to increased arousal frequency and, therefore, reduced survival. We predicted that hibernaculum conditions that reduce EWL (cold temperatures, high relative humidity) would increase our modeled survival rates. We also predicted that model parameters that influence EWL (surface area, area-specific rate of EWL), would have greater effects on modeled survival rates compared to other parameters.

Building on equations developed by Thomas et al. [1], Humphries et al. [24], and Hayman et al. [25], we included the effects of hibernaculum microclimate on fungal growth, EWL, torpor bout duration, and total fat loss. We parameterized the model using morphometric and physiological characteristics collected from *M. lucifugus* captured in the field. We validated the modified model components using a variety of data sources, determined the most influential parameters using a sensitivity analysis, and predicted fat loss over a range of hibernaculum conditions for both healthy and *P. destructans-*infected bats. Finally, in the context of winter duration, we inferred the impact of WNS on survival by comparing pre-hibernation fat stores to fat loss estimated by the energy expenditure model.

## Methods

### Ethics statement

All procedures were approved by the Texas Tech University Institutional Animal Care and Use Committee (protocol 16031-05) and followed guidelines of the Guide for the Care and Use of Laboratory Animals. We obtained proper permits from the Montana Department of Fish, Wildlife & Parks (permits 2016-104, 2017-018, and 2018-008).

### Study species

*M. lucifugus* is a common insectivorous bat species found across most of North America [26]. The hibernation behavior of *M. lucifugus* is well-studied. During the winter, *M. lucifugus* hibernate in caves and abandoned mines, often in large colonies [26]. Most hibernacula have stable microclimates, with high humidity (≥ 90 % RH) and temperatures ranging from 2 to 8 °C [8,27,28]. Many energetic models have determined energy expenditure during hibernation in response to microclimate selection, sex, and location [1,8,24,29–35]. Energetic models have been used to predict energy expenditure from WNS with alterations to arousal frequency [21,32]. *M. lucifugus* is also one of the most studied species in terms of WNS impacts. Since the discovery of WNS in 2006, millions of *M. lucifugus* have died across the species’ range and have faced upwards of 99% mortality rates [28,36,37]. Populations across eastern and mid-western North America affected by WNS remain at severely reduced population sizes and reduced population growth rates [36,38,39].

### Field data collection for model parameters

We captured *M. lucifugus* during the pre-hibernation (September-November) swarming and mid-hibernation (January-February) periods from 2016-2018 at a cave in central Montana. We used mist nets placed at the cave entrance to capture bats during swarming and hand-captured bats from hibernaculum walls during mid-hibernation. We transported bats in cloth bags to a mobile laboratory at the field site location for morphometric measurements. We weighed each bat (± 0.1 g) and used quantitative magnetic resonance (Echo-MRI-B, Echo Medical Systems, Houston, TX) to measure fat mass and lean mass [40]. We measured torpid metabolic rate (TMR) and EWL using open-flow respirometry at 2, 5, 8, and 10 °C (Supplementary Materials; [41]). We calculated the mean body mass, fat mass, and lean mass across all fall field seasons and the mean of mass-specific TMR across both seasons among all individuals across to use as parameters in the hibernation model (Table 1).

**Table 1.** Parameters for the energetics model for the little brown bat (*Myotis lucifugus*), their units, and the reference.

| Parameter Name | Parameter | Value | Units | Reference |
| --- | --- | --- | --- | --- |
| Basal metabolic rate | BMR | 2.6 | ml O<sub>2</sub> g<sup>-1</sup> h<sup>-1</sup> | Calculated from [55] |
| Minimum torpid metabolic rate | TMR<sub>min</sub> | 0.14 | ml O<sub>2</sub> g<sup>-1</sup> h<sup>-1</sup> | Measured in this study |
| Lower defended temperature during torpor | T<sub>tor - min</sub> | 2 | °C | [48–50] |
| Lower critical temperature | T<sub>lc</sub> | 32 | °C | [48–50] |
| Euthermic body temperature | T<sub>eu</sub> | 37 | °C | [1,48–50] |
| Change in torpid metabolism | Q<sub>10</sub> | $1.6 + 0.26 T_a - 0.006 T_a^2$ | - | [48] |
| Torpid thermal conductance | C<sub>t</sub> | 0.20 | ml O<sub>2</sub> g<sup>-1</sup> °C<sup>-1</sup> h<sup>-1</sup> | Calculated from [73] |
| Euthermic thermal conductance | C<sub>eu</sub> | 0.26 | ml O<sub>2</sub> g<sup>-1</sup> °C<sup>-1</sup> h<sup>-1</sup> | Calculated from [27] |
| Wing surface area | SA<sub>wing</sub> | 19.68 | cm<sup>2</sup> | Calculated in this study |
| Body surface area | SA<sub>body</sub> | 39.26 | cm<sup>2</sup> | Calculated from [53] |
| Area-specific rate of evaporative water loss for wing | rEWL<sub>wing</sub> | 0.33 | mg hr<sup>-1</sup> ΔWVP<sup>-1</sup> cm<sup>-2</sup> | Calculated in this study |
| Area-specific rate of evaporative water loss for body | rEWL<sub>body</sub> | 0.10 | mg hr<sup>-1</sup> ΔWVP<sup>-1</sup> cm<sup>-2</sup> | Calculated in this study |
| Time in euthermia per arousal | t<sub>eu</sub> | 1.10 | h | [31,74,75] |
| Maximum time in torpor | t<sub>tor-max</sub> | 1300 | h | [51] |
| Specific heat of tissue | S | 0.173 | ml O<sub>2</sub> g<sup>-1</sup> °C<sup>-1</sup> | [14] |
| Rewarming rate | WR | 0.80 | °C min<sup>-1</sup> | [44,58,76,77] |
| Body mass | M<sub>b</sub> | 7.80 | g | Measured in this study |
| Proportion of lean mass | pLean | 0.58 | g | Measured in this study |
| Proportion of fat mass | pFat | 0.26 | g | Measured in this study |
| Proportion of body water threshold | pMass | 0.027 | mg | Calculated in this study |
| Humidity-dependent fungal growth parameter | μ<sub>1</sub> | 1.51 x 10<sup>-4</sup> | - | [25] |
| Humidity-dependent fungal growth parameter | μ<sub>2</sub> | -9.92 x 10<sup>-3</sup> | - | [25] |
| Temperature-dependent fungal growth parameter | β<sub>1</sub> | 1.15 x 10<sup>-3</sup> | - | [25] |
| Temperature-dependent fungal growth parameter | β<sub>2</sub> | 0.27 | - | [25] |

We measured hibernaculum temperature and relative humidity over each hibernation period using HOBO (Model U23-001, ± 0.001 °C, ± 0.001% RH, Onset Computer Corporation) and iButton (temperature only; Model DS1921Z-F5, ± 0.05 °C, Maxim Integrated Products) dataloggers. We placed four HOBO and ten iButton dataloggers throughout the hibernaculum in the fall and recorded conditions at 3 h intervals. We determined the main winter roosting location from personal communication with U.S. Forest Service and Montana Department of Fish, Wildlife, and Parks personnel. We placed two HOBO loggers in the main roost, a large cathedral room at the back of the cave (one logger at the far end, one at the entrance), one within 3 m of the entrance of the cave, and one attached to a tree immediately outside the cave entrance (< 10 m). We spaced the iButtons evenly throughout the cave system from the entrance to the cathedral room. We suspended HOBO loggers with copper wire and used pantyhose to attach each iButton to a projected rock to suspend the logger in the air column. We collected loggers from the hibernaculum in the spring of each year.

We estimated winter duration for central Montana by acoustically monitoring bat activity at the entrance to the cave (Anabat Roost Logger RL1, Titley Scientific). The acoustic logger operated between 30 min before sunset and 30 min following sunrise. We used AnaLookW software (v4.3) to digitize calls and count the number of bat passes per day [42]. We were not interested in species-specific calls, but rather use the calls as an index of winter duration so we counted passes that contained calls of *Myotis* species (minimum frequency [f_min_] > 30 kHz) to filter out noise [43]. We were also not interested in the number of individual bats passing the detector, but rather if there was general activity outside the cave; we thus used a threshold of 50 passes day^−1^, defining the lower end of the 95% of bat counts, to determine the onset and end of the hibernation period [43].

### Incorporating evaporative water loss into the hibernation energetics model

We revised the hibernation energetics model first described by Thomas et al. [1] and Humphries [24], and then modified to include fungal growth by Hayman et al. [25]. The model estimates the amount of fat consumed during hibernation as a summation of the energy expended during multiple torpor-arousal bouts across a winter period (full model presented in Supplementary Materials). We derived estimates of the energy required during torpor (E_tor_), euthermia (E_eu_), and the warming (E_warm_) and cooling (E_cool_) periods between torpid and euthermic temperatures. We estimated the period of each arousal spent within euthermia from literature (Table 1) and the time to warm and cool were calculated given published warming and cooling rates, respectively [44].

We incorporated a mechanistic link between EWL and torpor bout duration. We estimated torpor bout duration (t_tor_) in two ways: 1) as a function of torpid metabolic rate in response to ambient temperature (T_a_) as described in Hayman et al. [25], and 2) as a function of EWL. Our revised model uses the shorter of the two estimates given hibernaculum conditions (Supplementary Figure S1), either arousing as a consequence of EWL or TMR, whichever comes first. By including both calculations in our estimates of torpor bout duration, we considered both the effect of EWL and metabolism on torpor physiology [45,46].

To estimate torpor bout duration as a function of metabolic rate (t_torTMR_), we modified the existing equations developed by Hayman et al. [25] that scale maximum possible time in torpor (t_torMax_) by the effects of metabolic rate given T_a_:

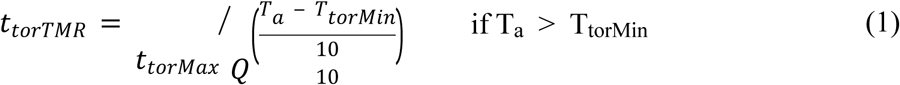

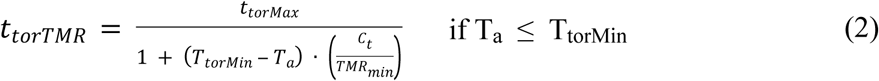

where Q_10_ is the change in metabolism with a 10°C change in temperature [47], T_torMin_ is the minimum defended T_b_ in torpor, TMR_min_ is the associated metabolic rate at T_torMin_, and C_t_ is the thermal conductance during torpor. Minimum defended T_b_ [48–50] and the maximum time in torpor (t_torMax_) were estimated from literature [51], and minimum torpid metabolic rate and thermal conductance were measured in the field using respirometry (Supplementary Materials).

To calculate torpor bout duration as a function of EWL (t_torEWL_), we assumed bats arouse when the total body water pool was depleted to a threshold [5]. The hourly rate of total EWL (mg H_2_O h^−1^) is comprised of both cutaneous and respiratory rates of EWL and is dependent on the water vapor pressure deficit between the bat and the surrounding environment. The hourly rate of cutaneous evaporative water loss (CEWL; mg H_2_O h^−1^) is a function of the difference between water vapor pressure at the surface of the bat and the environment (ΔWVP):

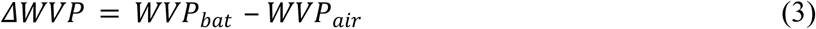

where WVP_bat_ is the water vapor pressure at the skin surface and WVP_air_ is the water vapor pressure of the surrounding air (both in kPa). We assumed WVP_bat_ was at saturation, which can be calculated as:

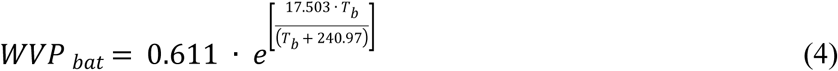

where T_b_ is the body temperature of the bat in torpor [52]. We then calculated WVP_air_ at T_a_ and given relative humidity. We modeled cutaneous EWL as a function of ΔWVP and the area-specific rate of EWL from bodily tissue (rEWL; mg H_2_O h^−1^ cm^−2^ per ΔWVP^−1^) across the surface area (SA; cm^2^) of the bat:

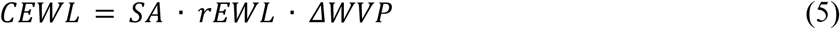

We used a surface area scaling equation [53] to calculate body surface area (SA_body_) and photos of bat wings to estimate the total surface area of the wings and tail (SA_wing_; Supplementary Materials). Assuming that a furred body and naked wing have biophysical differences that would affect cutaneous EWL, we used different values of the area-specific rate of EWL for the body (rEWL_body_) and wing (rEWL_wing_), estimated from respirometry (Supplementary Materials). Therefore, we rewrote Equation 5 as:

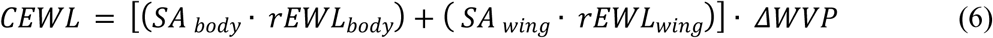

Respiratory EWL (REWL; mg H_2_O h^−1^) is a function of the saturation deficit between inspired and expired air. We assumed that inspired air is at T_a_ and is expired as saturated air at torpid T_b_ [5]. Therefore, we calculated respiratory EWL as:

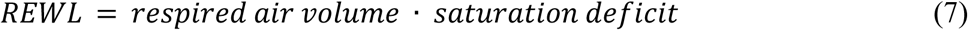

The volume of air that a bat breathes per hour was calculated as a function of the respiration rate of oxygen (i.e. TMR_min_) in ml O_2_ g^−1^ h^−1^ and body mass:

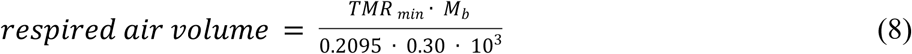

assuming the fractional concentration of oxygen in air is 0.2095 and that oxygen extraction efficiency is 30% [5]. Using the ideal gas law [52], we converted the water vapor pressure deficit (*ΔWVP*; Equation 3) from kPa to mg L^−1^ to determine the saturation deficit.

We validated the rate of total EWL (cutaneous EWL and respiratory EWL) by comparing modeled EWL (from Equations 5 and 7) to measured EWL from each individual during our respirometry procedures. We used individual body mass (Equations 5-6), metabolic rate (Equations 7-8), area-specific rate of EWL (Equations 5-6), and predicted surface area given body mass (Equations 5-6). We modeled the hourly rate of total EWL given the measured T_a_ and WVP experienced by each individual. We used linear regression to compare modeled EWL to measured EWL rates, assuming that if the model was accurate, the slope of the relationship should be equal to 1.

Given total EWL, we calculated torpor bout duration (t_torEWL_) based on the reduction of the total body water pool, setting the threshold at 2.7% of lean mass (assuming no body water in fat stores):

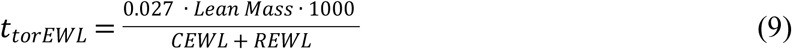

### Including the effects of fungal growth on hibernation

We further adjusted the hibernation model by including a link between fungal growth and reduced torpor bout duration through an increase in both metabolic rate and EWL (modifying Equations 1-2, 9). We first altered the estimation of torpor bout duration from T_a_ (t_torTMR_; Equations 1 and 2) by scaling t_torTMR_ by the proportion the bat wing surface affected by the fungus. When fungal growth > 0, t_torTMR_ was calculated as:

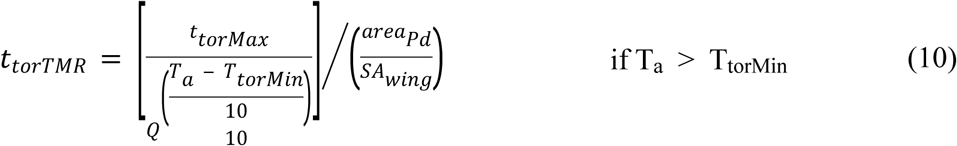

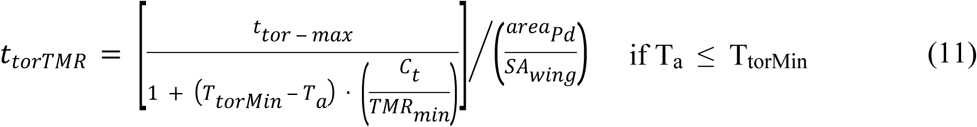

where *area*_*Pd*_ is the area (cm^2^) of fungal growth calculated as a function of T_b_ and relative humidity given equations from Hayman et al. [25].

We adjusted the calculation of torpor bout duration in response to EWL (t_torEWL_; Equation 9) by increasing CEWL and REWL as a function of fungal growth. We used data from McGuire et al. [20], who directly measured an increase in TMR and EWL in *M. lucifugus* infected with *P. destructans* (Supplementary Materials). We increased CEWL by including a linear increase to the rate of EWL of bat wings (rEWL_wing_) in response to the proportion the bat wing surface affected by the fungus (from Equation 6):

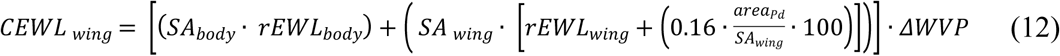

where 0.16 is the rate of increase in rEWL_wing_, given the proportion the bat wing surface affected by the fungus, determined from data presented in McGuire et al. [20] (Supplementary Materials). REWL also is hypothesized to increase in response to fungal growth with an increase in TMR; we included this linear increase by adjusting Equation 8:

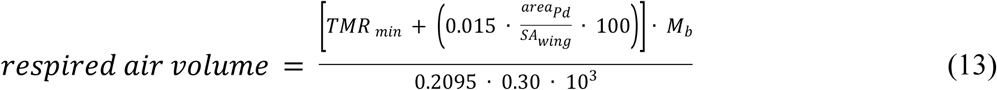

where 0.015 is the linear increase of torpid metabolic rate given the proportion the bat wing surface affected by the fungus (Supplementary Materials).

To validate the adjustment to the estimation of torpor bout duration in response to fungal growth (Equations 10-11,13), we used an independent dataset of skin temperature measurements from a captive hibernation study by McGuire et al. [54]. Skin temperature data were measured from thirteen *M. lucifugus* infected with *P. destructans* prior to hibernating in a controlled environment (T_a_ = 7 °C, relative humidity = 98%). Using methodology from Jonasson and Willis [31], we defined torpor and arousal periods based on cut-off temperatures and calculated the total time in each hibernation phase. We then estimated torpor bout duration (Equations 10-13) at each measured torpor bout from each individual given individual morphometric parameters (initial body mass, predicted surface area). We estimated TMR from body mass and T_b_ [55] and allowed for variation in lean mass (to determine threshold of body water) by sampling from a normal distribution with mean and standard deviation from our capture data. We predicted fungal growth area at each torpor bout given the time since inoculation and equations 2-4 in Hayman et al [25]. We then used a linear model to compare modeled torpor bout duration to measured torpor bout duration, assuming that if the prediction was accurate, the slope of the relationship should be equal to 1. To determine if including EWL improved our description of torpor expression, we also predicted torpor bout duration without the contribution of EWL using only Equations 10-11. We then compared these predictions to measured bout duration to determine model accuracy. Finally, we compared the R^2^ values of both fitted relationships to determine which model had better precision in predicting torpor bout duration.

### *Estimation of total fat loss and survival for* M. lucifugus *in Montana*

Using our modified hibernation model and model parameters obtained from our field captures and literature (Table 1), we estimated time until total fat exhaustion for *M. lucifugus* over the range of hibernaculum microclimate conditions measured at our field site. Torpor bout duration changes with body condition and fungal growth so we used differential equations to estimate energy consumption over the winter. We assumed that bats require energy to arouse at the end of hibernation and to leave the hibernaculum in order to obtain food. Therefore, we included energy required to warm (E_warm_) and spend 24 h in euthermia (24 x E_eu_) at the end of winter hibernation. We used the lsoda function of the *deSolve* package, which allowed torpor bout duration to change over time given fungal growth, bat parameters (Table 1), and hibernaculum microclimate. We converted total energy consumed over time from ml O_2_ g^−1^ to the amount of fat expended (g) as:

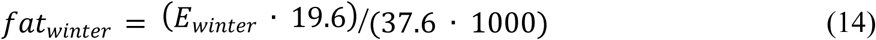

assuming that 1 ml O_2_ releases 19.6 J of energy and the energy content of fat is 37.6 J mg^−1^ [30]. We calculated time until fat exhaustion (t_fatEx_) as the time when total fat exhaustion (fat_winter_), became greater than mean fat stores measured during our fall field captures. Finally, we compared the estimated t_fatEx_ for both healthy and infected bats over the range of hibernaculum conditions to the duration of winter for central Montana estimated from our acoustic data. We assumed that mortality would occur if t_fatEx_ was less than winter duration; that is, the mean fat stores did not provide enough fat for a bat to survive through winter, as measured above.

We validated the entirety of the hibernation energetic model by comparing measured mass loss from 56 free-living hibernating *M. lucifugus* (Norquay and Willis, unpublished data, but see [56] for description of capture methodology and locations) to predicted fat loss from our model. We used this dataset because data from captive animals may not accurately reflect field conditions of free-living animals. We used individuals in which mass was measured during both swarming (August-September) and emergence (April-May). We estimated fat loss using the bioenergetic model for the time between swarming and emergence capture dates, given the hibernaculum conditions where each bat was captured [57,58]. We took the mean and standard deviation of T_a_ and water vapor pressure of each capture location and sampled random values from a normal distribution for each individual. We estimated TMR from body mass and T_b_ [55] and allowed for variation in lean mass by sampling from a normal distribution set at the mean and standard deviation from our capture data. We compared estimated fat loss with measured mass loss (assuming all mass change is due to fat loss) using linear regression, assuming if the two values were the same, the slope of the relationship would be no different than 1. We also predicted fat loss for the validation dataset given the hibernation model without the inclusion of EWL; more specifically, we only included Equations 1-2 in our calculations of torpor bout duration. We compared these predictions to measured mass loss and determined both model accuracy (slope = 1) and precision (R^2^) to compare against our modified model including EWL.

Following Hayman et al. [25], we used a multi-parameter sensitivity analysis to assess the impact of each parameter on estimations of time until mortality. Using Latin hypercube sampling in R package *lhs*, we created 100 random parameter sets sampled from a uniform distribution of potential values ranging from 10% lower or higher than the default value (Table 1). By constraining the minimum and maximum values of the parameters, and including a joint distribution within the Latin hypercube sampling, we considered the potential for correlations between parameters. We determined the relative importance of each variable by comparing partial rank correlation coefficients (PRCC) values. Positive PRCC values indicate an increase in the model output with an increase in the parameter value, while negative PRCC values indicate a decrease in the model output with an increase in parameter value [25].

## Results

We captured 219 *M. lucifugus* over the capture periods of 2016-2018 (176 during fall, 43 during winter; Table 2). There was minimal variation in hibernaculum microclimate measured by the HOBO and iButton loggers within the hibernaculum (temperature: mean = 4.80 ± 0.60 °C, range = 2.77 − 5.68 °C; water vapor pressure deficit: mean = 0.11 ± 0.26 kPa, range = 0.00 – 2.57 kPa) across winters (Figure 1). We found all bats roosting in the cathedral room, where hibernaculum microclimate was stable throughout the winter (Ta = 4.8 °C, RH = 100%). Activity decreased < 50 passes day^−1^ by mid-October (mean date among years 14 October) and increased beyond 50 passes day^−1^ by mid-April (mean date among years 13 April). We therefore concluded that hibernation duration in central Montana was 181 days.

**Table 2.**
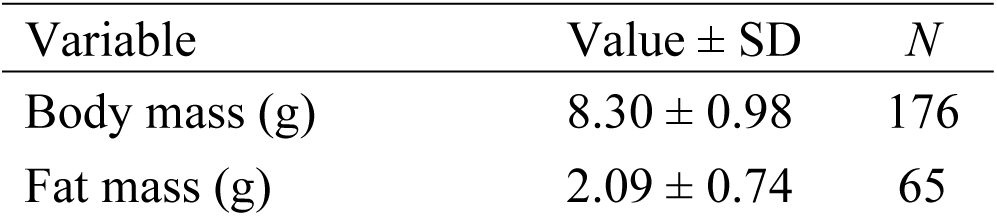
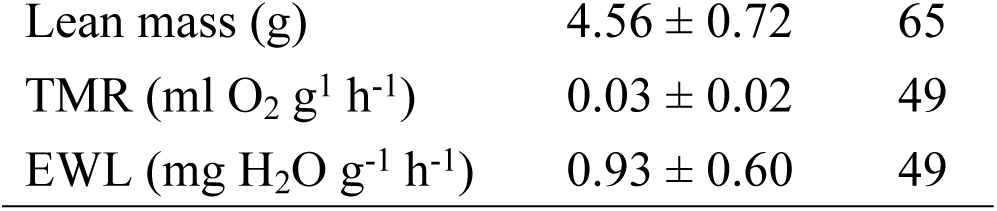
Morphometric and physiological data measured from *Myotis lucifugus* captured at a hibernaculum in central Montana. *N* = sample size, TMR: mass-specific torpid metabolic rate, EWL: mass-specific evaporative water loss.

| Variable | Value ± SD | *N* |
| --- | --- | --- |
| Body mass (g) | $8.30 \pm 0.98$ | 176 |
| Fat mass (g) | $2.09 \pm 0.74$ | 65 |
| Lean mass (g) | $4.56 \pm 0.72$ | 65 |
| TMR (ml O<sub>2</sub> g<sup>-1</sup> h<sup>-1</sup>) | $0.03 \pm 0.02$ | 49 |
| EWL (mg H<sub>2</sub>O g<sup>-1</sup> h<sup>-1</sup>) | $0.93 \pm 0.60$ | 49 |

**Figure 1.**
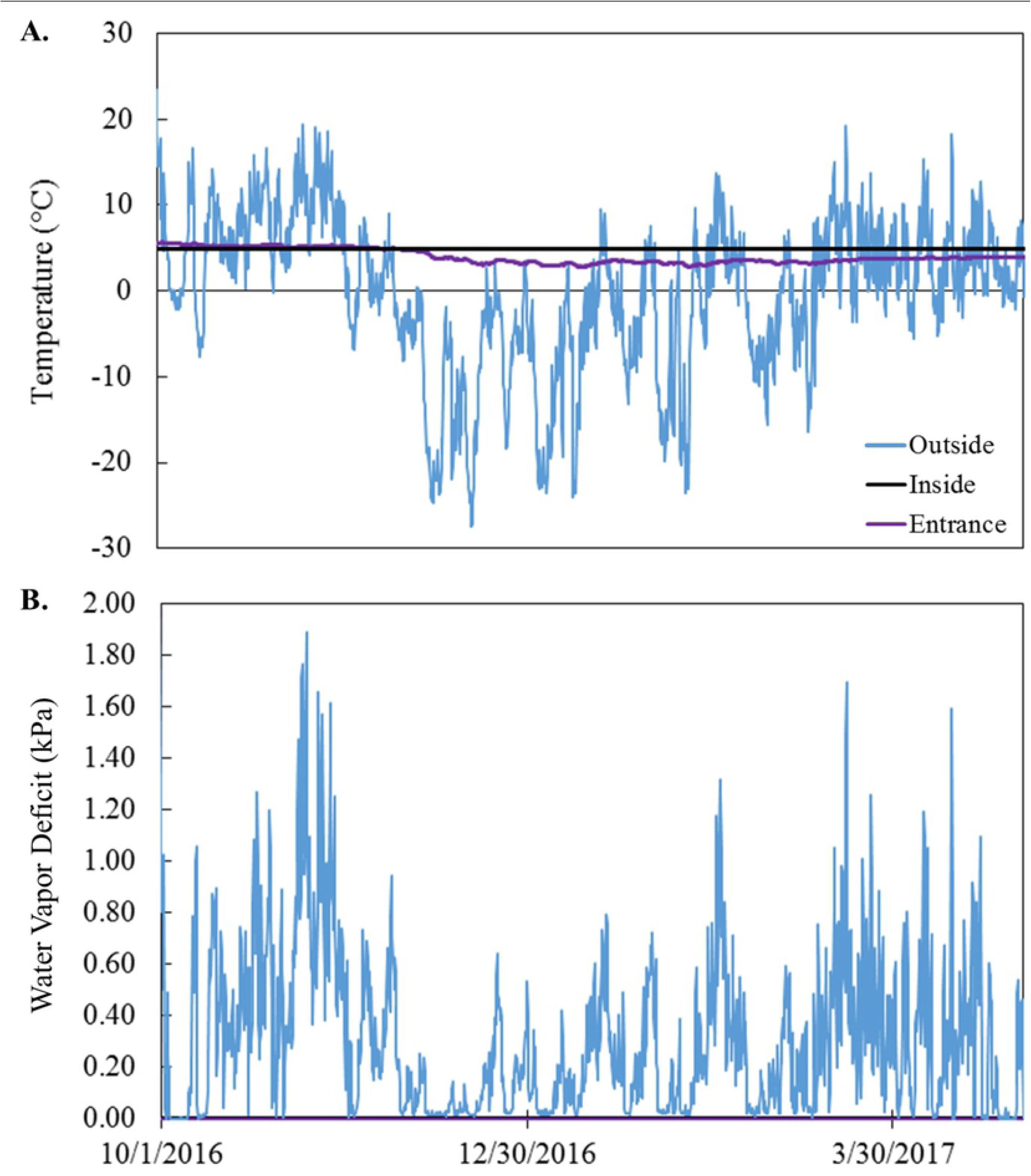
**(A)** Hibernaculum temperature (°C) and (**B**) water vapor pressure deficit (kPa) deep within the hibernaculum where *Myotis lucifugus* are found during hibernation (black line), at the entrance of the hibernaculum (purple line), and outside the hibernaculum entrance (blue line). Both the entrance (purple) and inside the hibernaculum (black) were at saturation the entire winter period.

Measured EWL from our respirometry procedures in dry air (0% relative humidity) ranged from 0.31 to 1.53 mg H_2_O h^−1^ g^−1^ (mean: 0.71 ± 0.25 mg H_2_O h^−1^ g^−1^) depending on temperature and individual. Our model accurately predicted EWL for *M. lucifugus* in Montana (*F*_1,61_ = 570.3, *p* < 0.001, *slope* = 0.97 [0.89, 1.06]; Figure 2a). Given the hibernaculum conditions measured at the roosting location (T_a_ = 4.8 °C, RH = 100%), we predicted EWL from *M. lucifugus* as 0.06 ± 0.40 mg H_2_O h^−1^ g^−1^ in healthy bats (Supplementary Figure S2). *P. destructans* had no impact on EWL early in infection, but by late hibernation had increased EWL to 2.19 mg H_2_O h^−1^ g^−1^ (Supplementary Figures S2).

**Figure 2.**
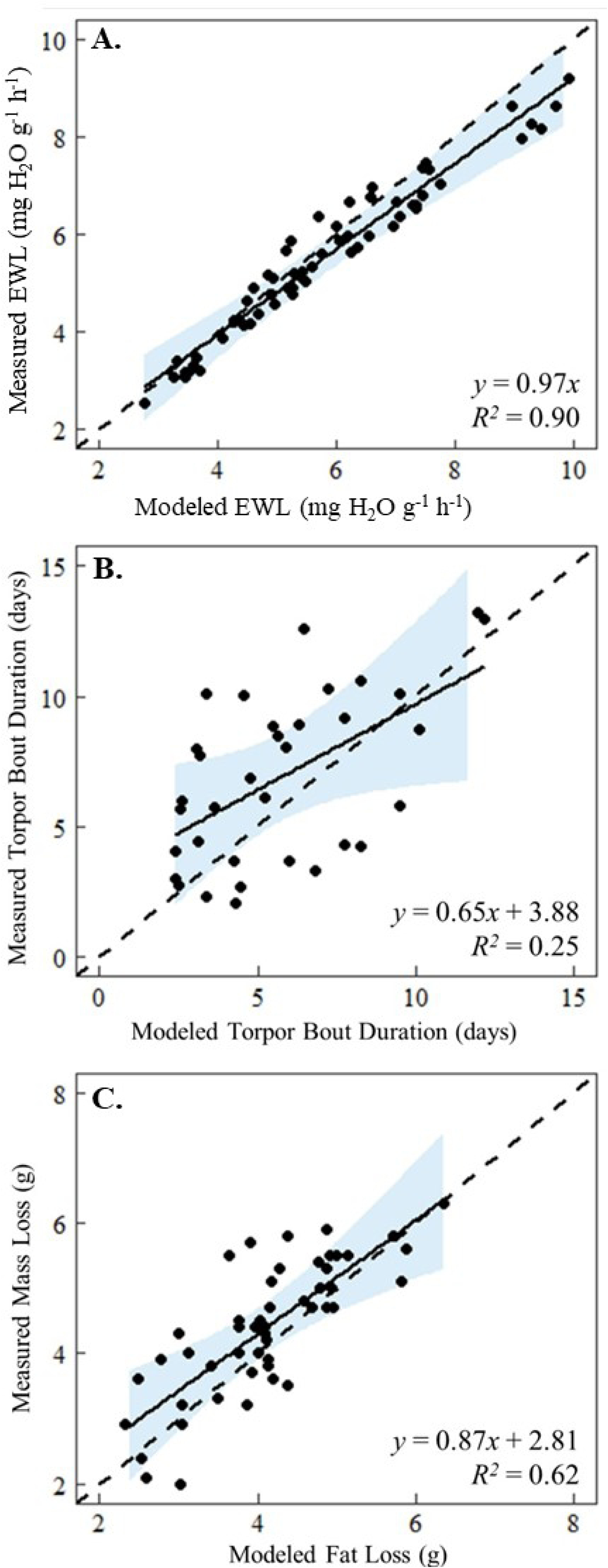
Comparison of measured and modeled (**A**) evaporative water loss (EWL), (**B**) torpor bout duration, and (**C**) fat loss in *Myotis lucifugus*. EWL and fat loss were measured/modeled in healthy bats, while torpor bout duration was measured/modeled in bats that were inoculated with *P. destructans*. Dashed lines represent one-to-one line and solid lines represent fitted relationship with 95% confidence intervals (shaded blue).

Our model accurately estimated torpor bout duration in captive bats infected with *P. destructans* (*F*_1,32_ = 18.64, *p* = 0.0001, *slope* = 0.65 [0.43, 1.16]; Figure 2b), but the estimates had a wide variance and lacked precision (only 25% of the variation in the data was explained by the model). Without the inclusion of EWL, however, the model did not accurately describe torpor bout duration (*F*_1,32_ = 0.40, *p* = 0.53, *slope* = −0.15 [-0.59, 0.30]) and did not describe variation in the data (R^2^ = 0.02). We therefore predicted torpor bout duration using our modified model including EWL. For healthy bats, torpor bouts lasted 16.10 ± 5.04 days within the microclimate conditions of the hibernaculum at the field site (range: 4.54 – 18.3 days; Supplementary Figure S3). Torpor bouts ranged from < 1 day to 18.3 days (mean: 6.20 ± 5.40 days) for bats infected with *P. destructans* (Supplementary Figure S3).

Our modified hibernation model accurately predicted mass loss in healthy wild bats (*F*_1,47_ = 74.38, *p* < 0.0001, *slope* = 0.87 [0.67, 1.07]; Figure 2c). Though there was a lack of individual metabolic rate and EWL data for the bats used in this validation procedure, our model still explained 62% of the variation in the dataset. Our model was also more precise than the hibernation model that lacked EWL, which was not accurate (*F*_1,47_ = 1.04, *p* = 0.84, *slope* = - 0.02 [-0.18, 0.15]) and described less than 1% of the variation in the data. Using the model with EWL, the mean time until total fat exhaustion for healthy *M. lucifugus* predicted in the hibernaculum microclimate conditions at our field site in Montana was 317.5 ± 105.50 days (Figure 3a) at a rate of 0.006 ± 0.002 g day^−1^. Bats were predicted to survive for over 360 days in the microclimate selected for roosting (T_a_ = 4.8 °C, RH = 100%). The shortest time until fat exhaustion (176 days) was at the warmest temperature available in the hibernaculum (5.5 °C) and lowest humidity (90%). Almost all other available microclimate conditions within the hibernaculum (2-5 °C and > 90% RH) result in predicted hibernation duration greater than winter duration (181 days).

**Figure 3.**
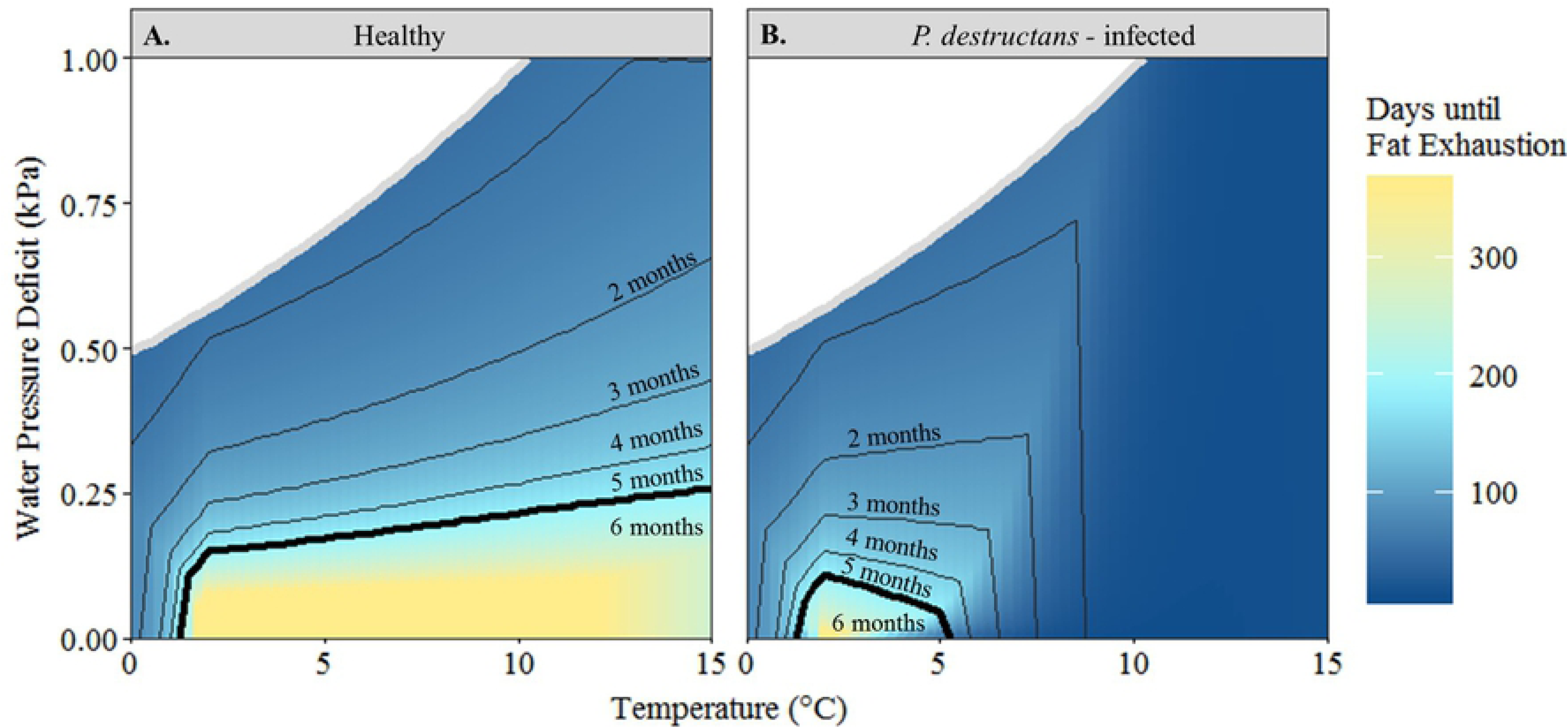
Predicted number of days until fat exhaustion for (**A**) healthy and (**B**) *P. destructans* - infected little brown bats (*Myotis lucifugus*) over a range of hibernaculum temperature (°C) and water vapor deficit (kPa) values. Contours represent hibernaculum conditions that allow survival for specific winter duration (in months); dark black contour indicates 6 months, the estimated hibernation duration at our study site in central Montana. White area bounded by grey line represents impossible parameters space for each temperature (e.g. at 2 °C, air is saturated at 0.50 kPa and cannot hold more water).

Within the hibernaculum conditions available at our field site, we predicted a higher and more variable rate of fat loss (range: 0.006 – 0.32 g day^−1^) for infected bats. In the specific hibernaculum conditions selected for roosting within these conditions, infected bats lost 0.01 ± 0.001 g day^−1^ at the beginning of hibernation (< 14 days) while the rate of fat loss increased to 0.03 ± 0.01 g day^−1^ at the end of hibernation (181 days). Almost all microclimate conditions available at our field site resulted in mortality for infected bats as time until fat exhaustion was less than hibernation duration (mean: 131.23 ± 38.40 days; Figure 3b). The only available microclimate conditions that permitted survival were at the lowest temperatures (2 – 3 °C) and highest humidity conditions (96 – 100%) but these locations were not selected by any healthy bats within this hibernaculum. Bats selected saturated environments that were within the temperature range that allowed fungal growth, resulting in increased energy expenditure and ultimately decreased time until total fat exhaustion.

Our sensitivity analysis revealed that fat loss was influenced by host-specific parameters, including body mass, the proportions of body mass comprised of fat and lean mass, and parameters that influenced EWL, including wing surface area and the area-specific rates of cutaneous EWL (Figure 4). Model parameters that were most influential to survival were physical traits that vary both within and among species. There was little effect of metabolic rate during torpor or euthermia, nor time spent euthermic.

**Figure 4.**
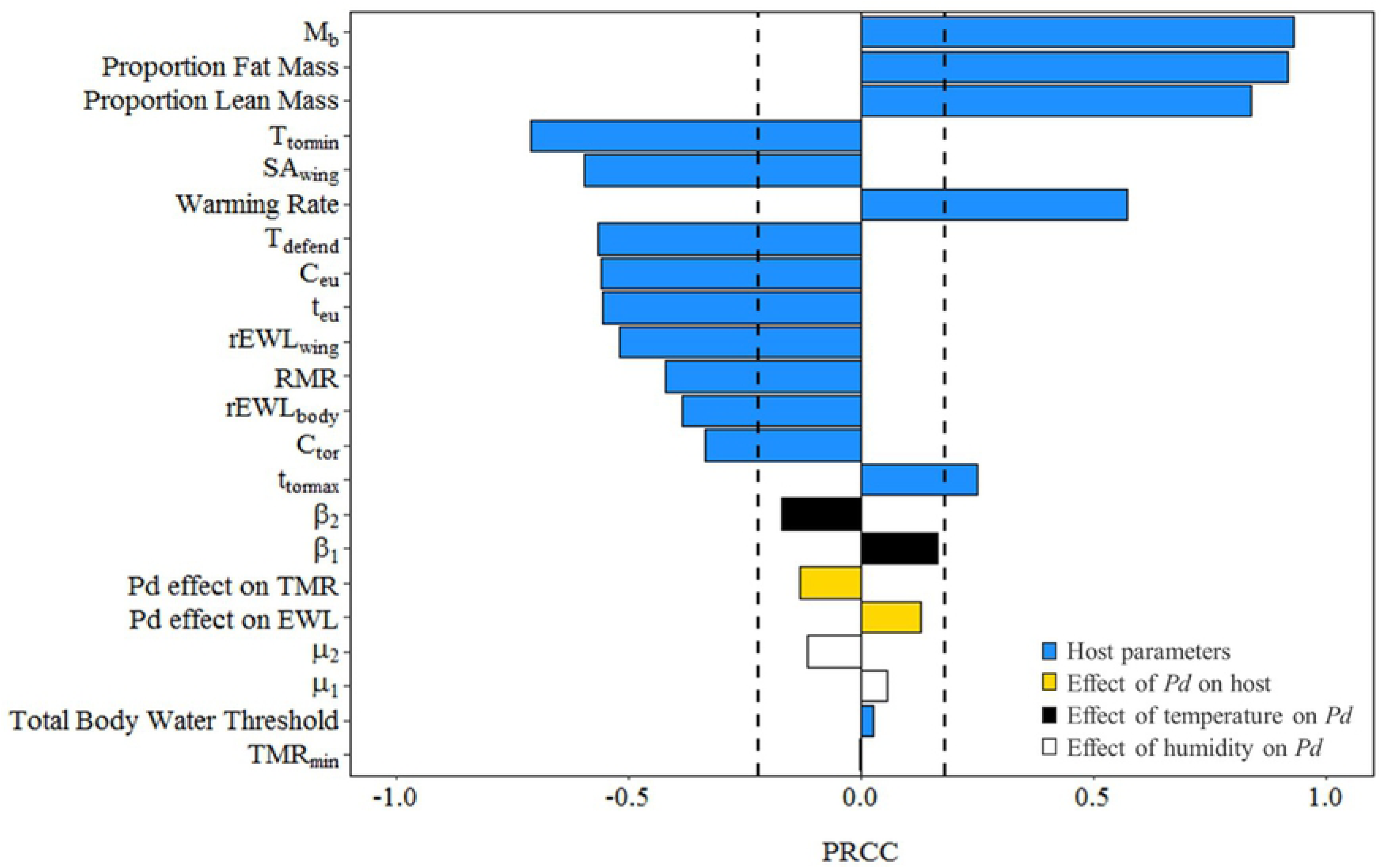
Sensitivity analyses for model calculating total fat exhaustion in hibernating bats infected with *P. destructans*. Dashed lines signify confidence intervals (α = 0.05). Positive PRCC values indicate an increase in predicted time until total fat exhaustion with an increase in parameter value; negative values indicate a decrease in predicted time until fat exhaustion with an increase in parameter value.

## Discussion

With the continued spread of WNS, it is imperative to understand the effects of hibernaculum microclimate (temperature and humidity) on fungal growth, host physiology, and winter survival. A model that includes effects of EWL on arousal frequencies within the study system of WNS, can improve understanding of the role of EWL on the evolution of hibernation and the interplay of host physiology with the environment. We showed that host parameters, particularly body mass, fat mass, and area-specific rate of EWL, were important drivers of torpor bout duration. Our results suggest that factors associated with EWL and arousal frequency are key elements for predicting the effects of WNS on hibernating bats.

Our modified hibernation bioenergetic model predicted torpor behavior similar to torpor-arousal behavior observed in wild *M. lucifugus* populations. For instance, Reeder et al. [59] measured torpor bouts of 16.32 ± 6.65 days and Czenze et al. [35] measured bouts of 16.2 ± 11.4 days in similar conditions where we predicted bouts of 16.1 ± 5.04 days in the Montana cave system (Supplementary Figure 2b). Observations of torpor behavior in WNS-affected bats corroborated our predictions of torpor bout duration (6.20 ± 5.40 days; Supplementary Figure 2b): wild populations of *M. lucifugus* remained in torpor for 7.93 ± 2.49 days [59], while captive populations averaged 6.48 ± 0.76 days [60]. Similarly, Reeder et al. [59] determined a negative relationship between wing damage due to fungal growth and torpor bout duration, which is aligned with how we incorporated the effect of infection in our model. Overall, the fidelity of the model implies that our prediction of torpor bout duration as a function of EWL is biologically relevant and representative of hibernation physiology and behavior.

We showed complete survival capacity (100% survival) in the entire microclimate space inhabited by healthy *M. lucifugus* in a cave system in central Montana (Figure 3a). Unfortunately, these hibernaculum temperatures and predicted torpor bout durations are comparable to hibernacula inhabited by highly impacted *M. lucifugus* populations in WNS-affected regions [28,35,39]. We predicted almost complete mortality (11% survival) for *M. lucifugus* within the current hibernaculum conditions in this cave system, in part because the high humidity selected by hibernating bats also results in high fungal growth [22,23]. Our predictions are consistent with population trends observed in WNS-affected regions in eastern North America, where similar hibernaculum microclimates have resulted in high mortality (80-98%) [39]. However, our model predicts a small window of microclimate space that would allow for survival, where cooler temperatures and moderate humidity reduce fungal growth, resulting in longer torpor bout duration and decreased arousal frequency (Figure 3b; Supplementary Figure S2). Our model predictions are consistent with observations of WNS-affected bats roosting in colder temperatures compared to unaffected bats [59,61]. These observations, in conjunction with our predictions, suggest that *M. lucifugus* within the Montana cave system would be highly impacted by WNS, but could potentially survive if individuals seek out cooler microclimates. Alternatively, if there are cooler microclimates available, those individuals that already prefer these conditions will survive while others will not. If microclimate preference is heritable, there is the potential for evolutionary rescue [62,63].

Our modified hibernation energetics model relies on measurements of the response of *P. destructans* to temperature from the lab and modeled response to relative humidity based on previous work [25]. Multiple studies reported diverse *P. destructans* responses to microclimate conditions [22,23,37,64,65], potentially due to subtle differences in laboratory procedures, and thus our predictions of fungal growth may not perfectly represent wild conditions. Additionally, parameters we used to estimate the increase in both metabolic rate and the rate of cutaneous EWL were derived from a single captive study [20]. However, results of studies of the effects of WNS on torpor patterns in wild and free-ranging bats are similar [59,66]. Additionally, our sensitivity analysis indicates that the predictions did not change significantly in response to changes in the temperature and humidity-dependent fungal growth rates or the increase to metabolic rate and EWL (Figure 4). Although future research into humidity-dependent fungal growth rate parameters on wild bats within natural conditions is warranted to increase our understanding of these dynamics, our predictions are consistent with the data currently available.

Evidence of at least some *M. lucifugus* populations with greater fat stores persisting post-WNS [67–71] corroborates our findings from our sensitivity analysis that fat mass is a major driver of WNS-survival (Figure 4). Large fat stores allow for increased arousal frequency associated with infection with *P. destructans* and extend the time until total fat exhaustion. Currently, we assess the costs of infection on the mean parameters – that is, body mass, fat mass, and lean mass represent the center of the distribution of morphometric characteristics if we assume a symmetrical distribution. Given evidence of the importance of body mass and fat, we would expect that some individuals from the Montana population would survive in the sampled hibernaculum if they had greater fat stores. It is therefore important that we further our research on the drivers of intra- and interspecific variation in overwintering survival from WNS.

The relationship between water vapor pressure and fungal growth indicates the potential for mitigation of WNS impacts if bats roost in microclimates below saturation – that is, infected individuals may trade-off water conservation with energy minimization. In healthy bats, maximum survival was at saturation (100% RH; Figure 3a). As saturation leads to negligible EWL [5], bats can remain in torpor longer before dehydration leads to arousal [46]; thus, roosting in saturated microclimates minimizes energetic costs. However, because *P. destructans* growth increases with water vapor pressure [23], infected bats had lower survival at saturation compared to less humid environments (Figures 3b). This hypothesis is supported by evidence of a relationship between increased population growth rate in multiple species and decreased relative humidity in regions post-WNS infection [39]; hibernacula with less than 90% relative humidity were the only microclimates to have population growth rates above zero, which aligns with our predictions of reduced survival at saturation.

Our model supports the role of EWL as a driver of periodic arousals in hibernation, and contributes to addressing one of the longest-standing questions in hibernation biology. It also showcases how interactions between host and pathogen physiology, and the environment can exacerbate or mitigate the costs of a disease. The relationship between EWL, fungal growth, and humidity suggests that bats found in some parts of western North America, where hibernacula are often drier than eastern hibernacula, may not be as impacted by WNS as eastern populations. Additionally, species and populations that inhabit more arid environments tend to have lower rates of EWL [72] due to adaptations to allow maintenance of water balance in sub-optimal conditions, and thus may not experience high WNS-related mortality [23]. The non-linear interplay of temperature, humidity, and behavior (selecting roosting conditions) needs further analysis, and our model provides a tool to address these questions. The model allows for species-specific parameterization and interspecific variation in morphometrics, physiology, and roosting habitats, suggesting that morphometric and physiological data from western bat species is needed. With this modified hibernation energetics model, we now have the tool to assess the potential impact of WNS on populations that have different hibernation behaviors than previously impacted species.

## Acknowledgments

We thank Quinn Fletcher and other members of the U. of Winnipeg bat lab for help with field data on mass loss of hibernating bats. We thank L. Hanauska-Brown for helping obtain permits and B. Maxwell and D. Bachen for site identification. We appreciate the field help from H. D. Bobbitt, D. Taylor, D. Jones, D. Crowley, E. Brandell, G. Botto, E. Lee, K. Smucker, and D. Bishop. This study was supported with equipment from Texas Tech University. This project has been funded in whole with Federal funds from the Department of Defense Strategic Environmental Research and Development Program, under Contract Number W912HQ-16-C-0015. RP was supported by DARPA D16AP00113, NSF DEB-1716698, NIH P20GM103474, and NIH P30GM110732. DTSH is supported by RDF-MAU1701. CKRW and KJON were supported by NSERC, Canada, USFWS, and Species at Risk Research Fund of Ontario.

## Author contributions

CGH wrote the manuscript and performed the analyses. CGH, NWF, KAS, KJON obtained field data. CGH, NWF, CRH, DTSH, SHO, and LPM developed the model. RKP, SHO, DTSH, and LPM acquired funding and designed the study. CGH, CRH, DTSH, NWF, LPM, RKP, CKRW, and SHO edited the manuscript.

## Data and materials availability

All data needed to evaluate the conclusions of the paper are available in the paper. Additional data or code related to this paper may be requested from the authors.

